# Visualization of the type III secretion mediated *Salmonella*-host cell interface using cryo-electron tomography

**DOI:** 10.1101/359166

**Authors:** Donghyun Park, Maria Lara-Tejero, M. Neal Waxham, Wenwei Li, Bo Hu, Jorge. E. Galán, Jun Liu

## Abstract

Many important gram-negative bacterial pathogens use highly sophisticated type III secretion systems (T3SSs) to establish complex host-pathogen interactions. Bacterial-host cell contact triggers the activation of the T3SS and the subsequent insertion of a translocon pore into the target cell membrane, which serves as a conduit for the passage of effector proteins. Therefore the initial interaction between T3SS-bearing bacteria and host cells is the critical step in the deployment of the protein secretion machine, yet this process remains poorly understood. Here, we use high-throughput cryo-electron tomography (cryo-ET) to visualize the T3SS-mediated *Salmonella*-host cell interface. Our analysis reveals the intact translocon at an unprecedented level of resolution, its deployment in the host cell membrane, and the establishment of an intimate association between the bacteria and the target cells, which is essential for effector translocation. Our studies provide critical data supporting the long postulated direct injection model for effector translocation.

## INTRODUCTION

Type III secretion systems (T3SSs) are widely utilized by many pathogenic or symbiotic Gram-negative bacteria to directly inject bacterially encoded effector proteins into eukaryotic host cells (Deng et al, 2017; Galán et al, 2014; Notti & Stebbins, 2016). The central element of the T3SS is the injectisome, a multiple-protein machine that mediates the selection and translocation of the effectors destined to travel this delivery pathway. The injectisome is highly conserved, both structurally and functionally, among different bacterial species including important pathogens such as *Salmonella, Yersinia, Shigella, Pseudomonas* and *Chlamydia* species. It consists of defined substructures such as the needle complex, the export apparatus, and the cytoplasmic sorting platform (Galán et al, 2014; Hu et al, 2017; Loquet et al, 2012; Schraidt & Marlovits, 2011; Worrall et al, 2016). The needle complex is composed of a membrane-anchored base, a protruding needle filament, and a tip complex at the distal end of the needle (Kubori et al, 1998; Schraidt et al, 2010; Schraidt & Marlovits, 2011; Worrall et al, 2016). The export apparatus, which is formed by several inner membrane proteins, functions as the conduit for substrate translocation across the bacterial inner membrane (Dietsche et al, 2016). The sorting platform is a large cytoplasmic multiple-protein complex that orderly selects and delivers the substrates to the export apparatus (Lara-Tejero et al, 2011).

In many bacterial species the activity of these protein injection machines is stimulated upon contact with the target eukaryotic cell plasma membrane, a process thought to be mediated by the tip complex (Barta et al, 2012; Blocker et al, 2008; Deane. JE et al, 2006; Ménard et al, 1994; Zierler & Galán, 1995). Host cell contact triggers a cascade of poorly understood events that lead to the deployment of the protein translocases onto the host cell plasma membrane to form a protein channel in the host cell membrane that mediates the passage of the effector proteins. In the case of the *Salmonella enterica* serovar Typhimurium (*S.* Typhimurium) T3SS encoded within its pathogenicity island 1, the protein translocases are SipB and SipC, which through a process that requires the tip protein SipD, are inserted in the host-cell plasma membrane to form the translocon channel (Collazo & Galán, 1997). Deployment of the translocon also results in the intimate association of the bacteria and the host cell, which is orchestrated by the protein translocases themselves (Lara-Tejero & Galán, 2009; Misselwitz et al, 2011). Despite the critical role of the translocases in intimate attachment and effector translocation, little is known about their structural organization when deployed in the host cell membrane, and previous attempts to visualize them did not provide distinct structural details. This paucity of information is due at least in part to the intrinsic difficulties of imaging bacterial/host cell interactions at high resolution. Here, we used bacterial minicells and cultured mammalian cells combined with high-throughput cryo-ET to study the initial interaction between *S*. Typhimurium and host cells. This experimental system allowed the visualization of the intact translocon deployed in the host cell plasma membrane, in contact with the tip-complex of the T3SS injectisome, at unprecedented resolution. This study provides new insights into the initial events of the T3SS-mediated bacteria-host cell interactions and highlights the potential of cryo-ET as a valuable tool for investigating the host cell-pathogen interface.

## RESULTS

### *in situ* structures of the T3SS injectisome in the presence or absence of protein translocases

An intrinsic property of many T3SSs is that their activity is stimulated by contact with the target host cell plasma membrane (Ménard et al, 1994; Zierler & Galán, 1995). This interaction results not only in the stimulation of secretion but also in the deployment of the protein translocases in the host cell membrane, a poorly understood process that is orchestrated by the tip complex of the injectisome’s needle filament. In the case of the *S*. Typhimurium SPI-1 T3SS the tip complex is thought to be composed of a single protein, SipD, which organizes as a pentamer at the tip of the needle filament (Rathinavelan et al, 2014). However, it has been previously proposed that in *Shigella* spp., in addition to IpaD, a homolog of SipD, the tip complex also contains IpaB, a homolog of SipB (Cheung et al, 2015). To get insight into the structural organization of the tip complex prior to bacterial contact with cultured ells, we compared the *in situ* structures of fully assembled injectisomes from minicells obtained from wild-type, ∆*sipB*, and ∆*sipD S.* Typhimurium strains (Fig. 1a-d, Extended Data Table 1). We found that injectisomes from wild-type or the ∆*sipB* strains were indistinguishable from one another. In contrast, injectisomes from a *∆sipD* strains exhibited a shorter needle (~45 nm) in comparison to the needle filaments of injectisomes from the wild-type or *∆sipB* strains (~50 nm). These observations suggest that SipD is the only structural component of the tip complex (Fig. 1e). To further explore this hypothesis, we examined by cryo-ET the injectisomes of minicells obtained from *S*. Typhimurium strains expressing FLAG-epitope-tagged versions of SipB, SipC, and SipD, after labeling with anti-FLAG antibodies (Fig. 1f-h). Only injectisomes from minicells obtained from the strain expressing SipD-FLAG showed the antibodies bound to the needle tip (Extended Data Table 2). This observation is consistent with the notion that, prior to cell contact, SipD is the main, and most likely only component of the tip-complex (Lara-Tejero & Galán, 2009).

**Figure 1.**
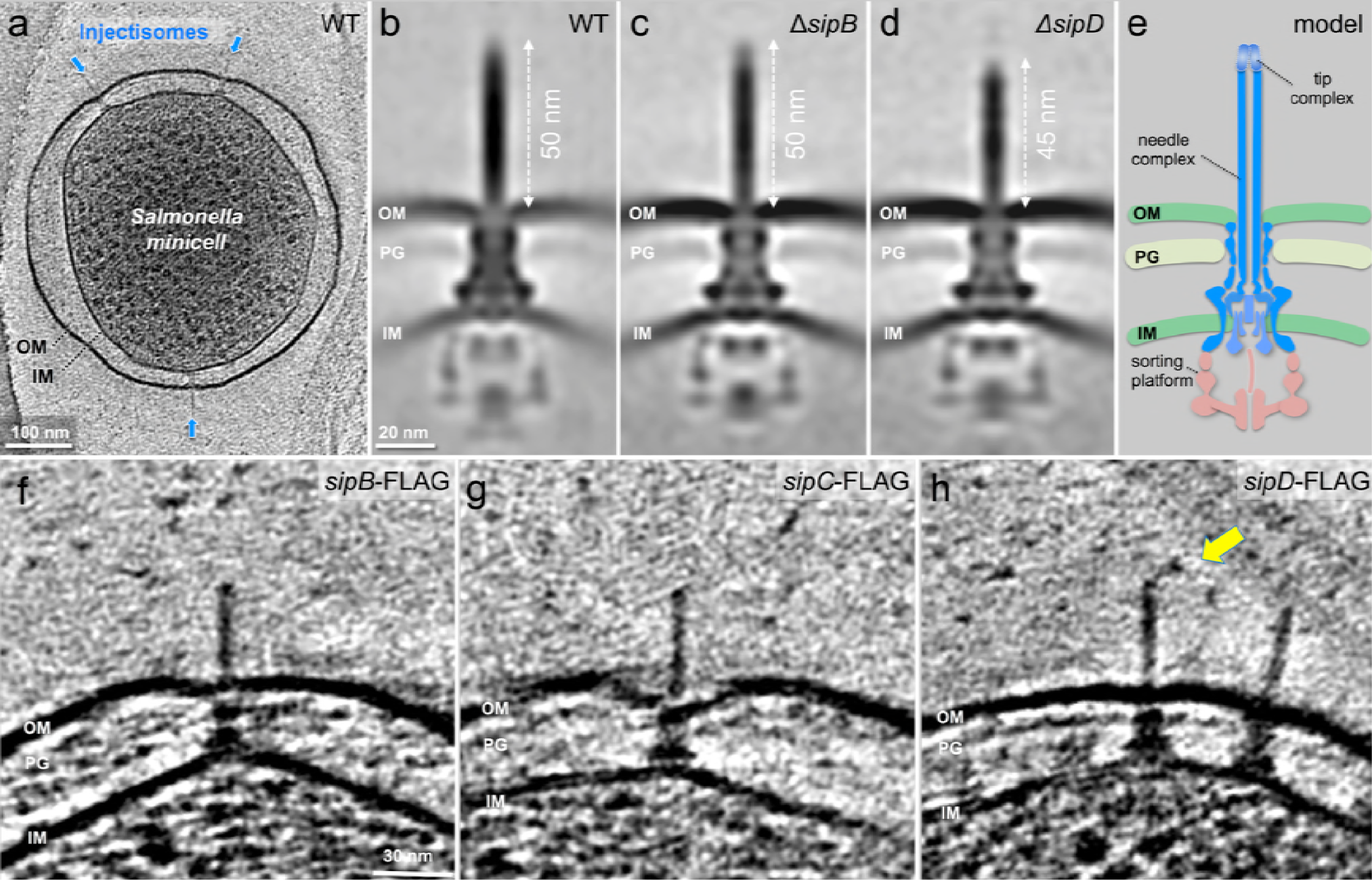
*In situ* structures of host-free *S.* Typhimurium T3SS injectisome in wild-type (WT), ∆*sipB*, and *∆sipD* minicells. (**a**) A central section of a tomogram showing *S.* Typhimurium minicell containing multiple injectisomes. (**b**-**d**) The central sections of sub-tomogram averages showing injectisomes of WT, ∆*sipB*, and ∆*sipD*, respectively. Outer membrane (OM), peptidoglycan (PG), and inner membrane (IM) of *S. Typhimurium* are annotated. (**e**) A schematic of the injectisome. (**f-h**) The central sections of tomograms showing injectisomes from strains expressing epitope-tagged (FLAG) SipB, SipC, and SipD, respectively. Yellow arrow indicates antibody bound to the epitope-tag.

### High-resolution imaging of the T3SS mediated *Salmonella*-host cell interface

It is well established that effector translocation through the T3SS requires an intimate association between the bacteria and the host cell (Grosdent et al, 2002). It has also been previously demonstrated that such intimate attachment requires an intact type III secretion machine, and in particular, the protein translocases, which most likely mediate such bacteria/host cell interaction (Lara-Tejero & Galán, 2009). Despite its central role in effector translocation, however, very little is known about the architecture of this specialized host/bacteria interface. This is largely because of the lack of amenable experimental approaches that would allow a detail view of this interface. Cryo-ET is uniquely suited to examine host/pathogen interactions at high resolution. However, sample thickness limits the utility of this approach. To get around this limitation we used bacterial minicells as a surrogate for whole bacteria since it has been previously shown that they are capable of assembling functional T3SS injectisomes that can deliver de novo synthesized T3SS substrates into cultured cells (Carleton et al, 2013). However, minicells are inefficient at triggering membrane ruffling, actin filament remodeling, and bacterial internalization due to inefficient partitioning of the effector proteins that trigger these responses. Consequently, while minicells are proficient at establishing a T3SS-mediated intimate association with cultured epithelial cells, they are inefficient at triggering their own internalization thus remaining firmly attached on the cell surface. These features make them ideally suited for high-resolution cryo-ET imaging. Therefore, we applied bacterial minicells obtained from wild-type *S.* Typhimurium onto cultured epithelial cells grown on cryo-EM grids. We found that the periphery of adherent cells is sufficiently thin (< 500 nm) for high-resolution imaging (Extended Data Fig. 1). We readily observed T3SS injectisomes at the interface between minicells and the plasma membrane of cultured epithelial cells (Fig. 2a-b). We found that in the presence of the injectisomes, the spacing between the surface of the *S*. Typhimurium minicells and the cultured-cell plasma membrane was ~50 nm, which matches the needle length of the injectisome imaged prior to their application to cultured cells (Extended Data Fig. 2a-f, m). The orientation of the injectisomes in the bacteria/target cell interface was perpendicular relative to the host PM, and the needle of the host-interacting injectisomes appeared straight (Fig. 2c). We also observed that the interaction of the injectisome and the target cell resulted in a noticeable inward bend of the PM (Extended Data Video 1). Consistent with this observation, the distance between the bacterial cell and the PM was shorter (~30 nm) than the distance observed in areas immediately adjacent to the injectisomes (Extended Data Fig. 2g-m). However, we did not observe any sign of penetration of the needle filament through the host cell plasma membrane as it has been previously proposed (Hoiczyk & Blobel, 2001). The length of the bacterial-envelope-embedded injectisome base substructure before (30.5 ± 2.3nm) and after (30.8 ± 2.2nm) the bacteria/target cell interactions remained unchanged (Extended Data Fig. 2m). This is in contrast to the *Chlamydia* T3SS, which has been reported to undergo significant conformational changes upon contact with host cells (Nans et al, 2015). The reasons for these differences is unclear and may either reflect intrinsic differences between these T3SS, or differences in the methodology used, which resulted in higher resolution of the visualized *S.* Typhimurium T3SS structures. Together, these observations indicate that (1) the interactions of the T3SS injectisome with the target cell results in the bending of the PM without penetration of the needle filament, and (2) upon contact with target cells the injectisome does not undergo conformational changes that could be seen at this level of resolution.

**Figure 2.**
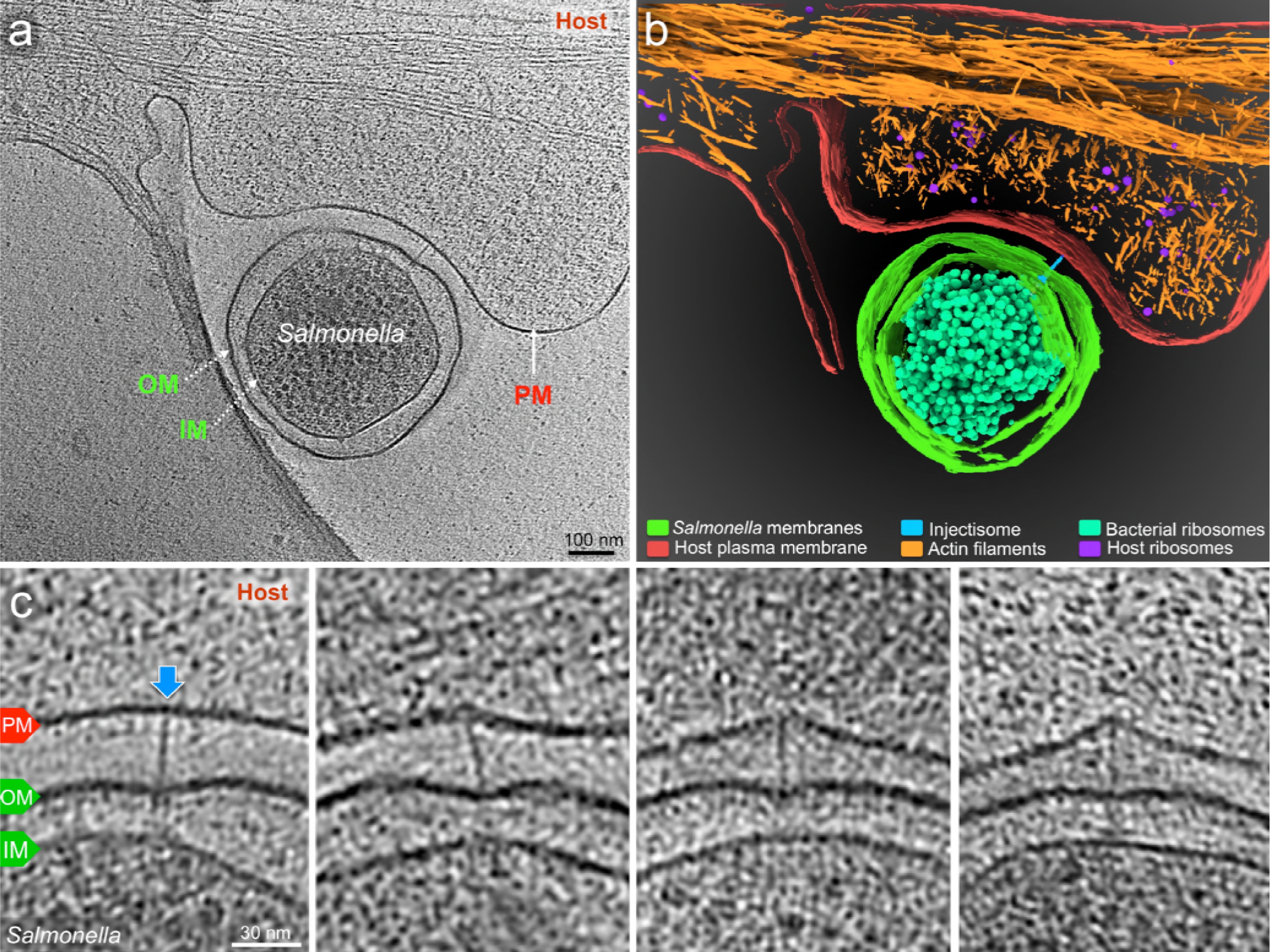
Visualization of the T3SS mediated *Salmonella-*Host interactions. **(a)** A central slice showing a *S.* Typhimurium minicell interacting with a host. Plasma membrane (PM) of HeLa cell, outer membrane (OM) and inner membrane (IM) of *S.* Typhimurium are annotated. **(b)** 3D rendering of the tomogram shown in (**a**). **(c)** Tomographic slices showing injectisomes interacting with the host PM. Blue arrows indicate needles attached to the host PM. Direction of the arrow represents the angle of needle perpendicular to the host PM.

### Visualization of the formation of the translocon in the target host cell membrane

The deployment of the translocon is an essential step in the T3SS-mediated delivery of effector proteins. However, very little information is available on both, the architecture of the assembled translocon, as well as the mechanisms leading to its deployment on the target cell. It is believed that the deployment process must be initiated by a sensing step most likely mediated by the tip complex (*i. e.* SipD), a step that must be followed by the subsequent secretion of the translocon components (*i. e.* SipB and SipC) destined to be inserted on the target eukaryotic cell PM. To capture the formation of the translocon, we analyzed over 600 injectisomes adjacent to the host PM. Classification of sub-tomograms depicting the region of the tip complex (Fig. 3a) showed the PM at various conformations and distances to the needle tip (Fig. 3b-i), which presumably represent intermediate steps prior to the deployment of the translocon and the resulting intimate attachment of the bacteria to the PM. After further alignment and classification of the injectisomes in intimate association with the PM, we obtained a distinct structure of the putative translocon in the host PM (Fig. 3j). Sub-tomogram averages of injectisomes from the *S.* Typhimurium translocase-deficient mutants ∆*sipB* or ∆*sipD* in close proximity to the target cell PM did not show this distinct structure, thus confirming that this density corresponds to the assembled translocon (Fig. 3k, l). To better visualize the translocon in 3D, we segmented the distinct translocon structure in the context of the host PM, the needle, and its tip complex (Fig. 3m, n). We found that the translocon has a thickness of 8 nm spanning the host PM and a diameter of 13.5 nm on its protruding portion (Fig. 3j). This size is substantially smaller than reported size of the translocon of enteropathogenic *E. coli* assembled from purified proteins *in vitro*, which was estimated to be 55-65 nm in diameter (Ide et al, 2001). One half of the translocon is embedded in the host PM, while the other half protrudes towards the host cytoplasm. In the middle of the protruded portion, we observed a hemispherical hole, which may represent the channel through which effectors make their way into the target cell plasma membrane. The presence of this structure is entirely consistent with the long-standing notion that the translocon forms a conduit through the host PM to facilitate the translocation of effectors (Mueller et al, 2008).

**Figure 3.**
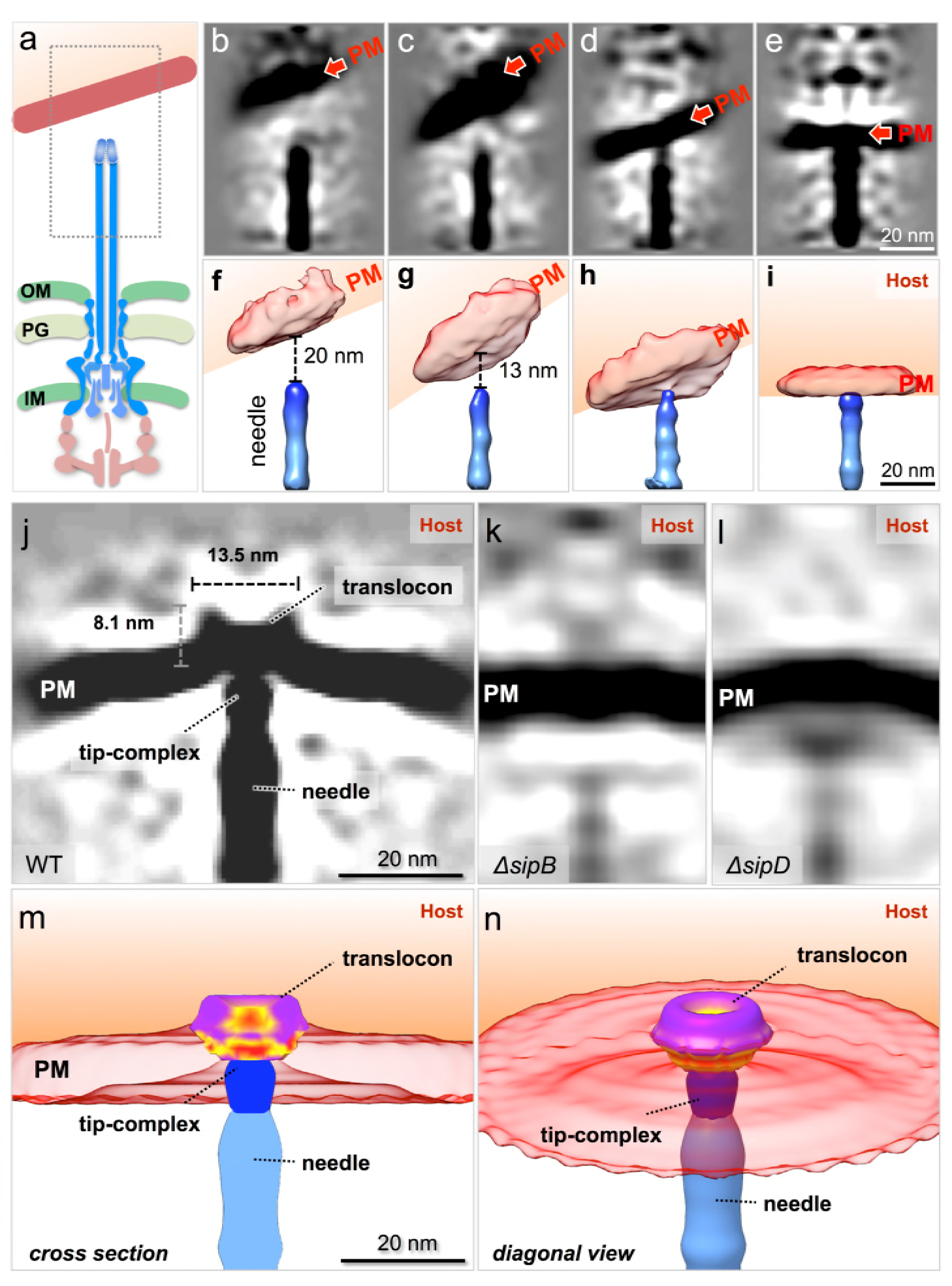
*In situ* structural analysis of the interface between the T3SS needle and the host membrane reveals a novel structure of the intact translocon. (**a**) A schematic representation of the *S.* Typhimurium injectisome with a box highlighting the area used for alignment and classification (**b-e**) Central sections and (**f-i**) 3-D surface views of class average structures showing different conformations of the needle - PM interaction. (**j-l**) Central sections of the sub-tomogram average structures of the interface between the host PM and the needle of WT, ∆*sipB*, and ∆*sipD*, respectively. Surface rendering of the structure in panel **j** in (**m**) a cross-section view and (**n**) a diagonal view.

Comparison of the arrangement of the injectisomes in relation to the target cell PM in wild-type and translocase-deficient strains revealed marked differences. In comparison to wild-type, bacterial cells obtained from translocase-deficient mutants showed a smaller proportion of injectisomes attached to the host PM (Fig. 4a). We also noticed that, unlike wild-type injectisomes, which most often appeared perpendicular to the target cell PM (Fig. 2c), the injectisomes from the translocase deficient mutant strains ∆*sipB*, ∆*sipD*, or ∆*sipBCD* appeared arranged at various angles relative to the PM. These observations are consistent with the fact that in the absence of the translocases, the injectisomes do not intimately attach to the target cell PM (Fig. 4b-j, Extended Data Fig. 3). These data also further support the notion that the distinct structure embedded in host membrane in close apposition to the T3SS injectisome needle tip is formed by the translocon.

**Figure 4.**
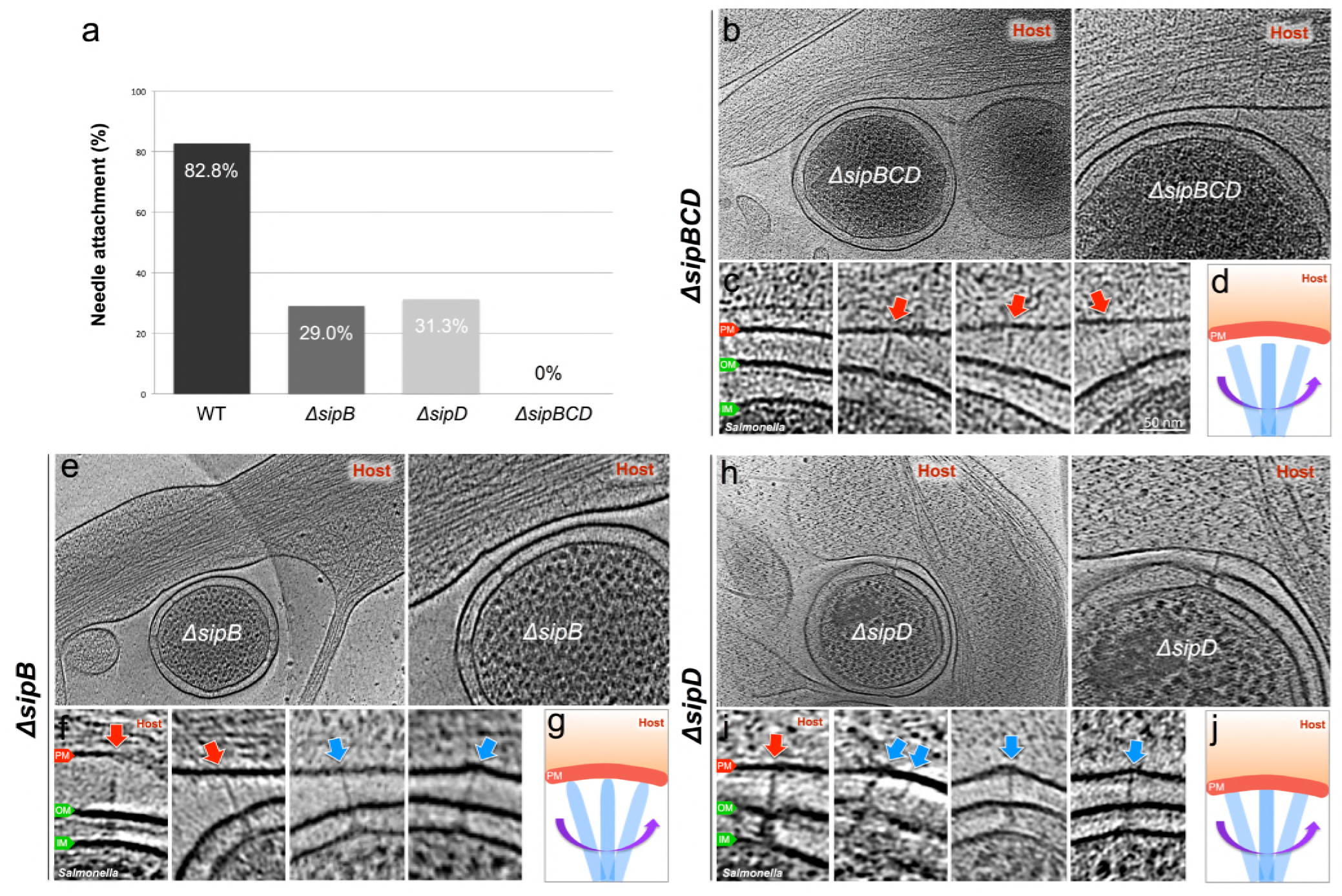
Deletion of the protein translocases disrupts the T3SS-dependent intimate attachment to the host PM, and the formation of the translocon. (**a**) Percentage of minicells attached to the host membrane via needle-membrane contact (**b, c, e, f, h, i**) Central slices from representative tomograms showing injectisomes interacting with the host PM. Blue arrows indicate needles attached to the host PM. Red arrows indicate unattached needles. Direction of the arrow represents the angle of needle perpendicular to the host PM. (**d, g, j**) Schematic models depicting needle-attachment patterns.

One of the striking features associated with the intimate T3SS mediated contact and the formation of the translocon is the target cell PM remodeling around the translocon-injectisome needle tip interface, appearing in a “tent-like” conformation (Fig. 2c, Extended Data Video 1). This feature is likely the result of the close association between the bacteria and the target cell presumably mediated not only by the T3SS but also by multiple additional adhesins encoded by *S*. Typhimurium. In fact, the distance of the bacteria OM and the target cell is shorter than the length of the needle itself, which results in the bending of the target cell PM and the “tent-like” conformation around the injectisome target cell PM interface. It is possible that this intimate association may facilitate the T3SS-mediated translocation of effector proteins (Fig. 5, Extended Data Video 2).

**Figure 5.**
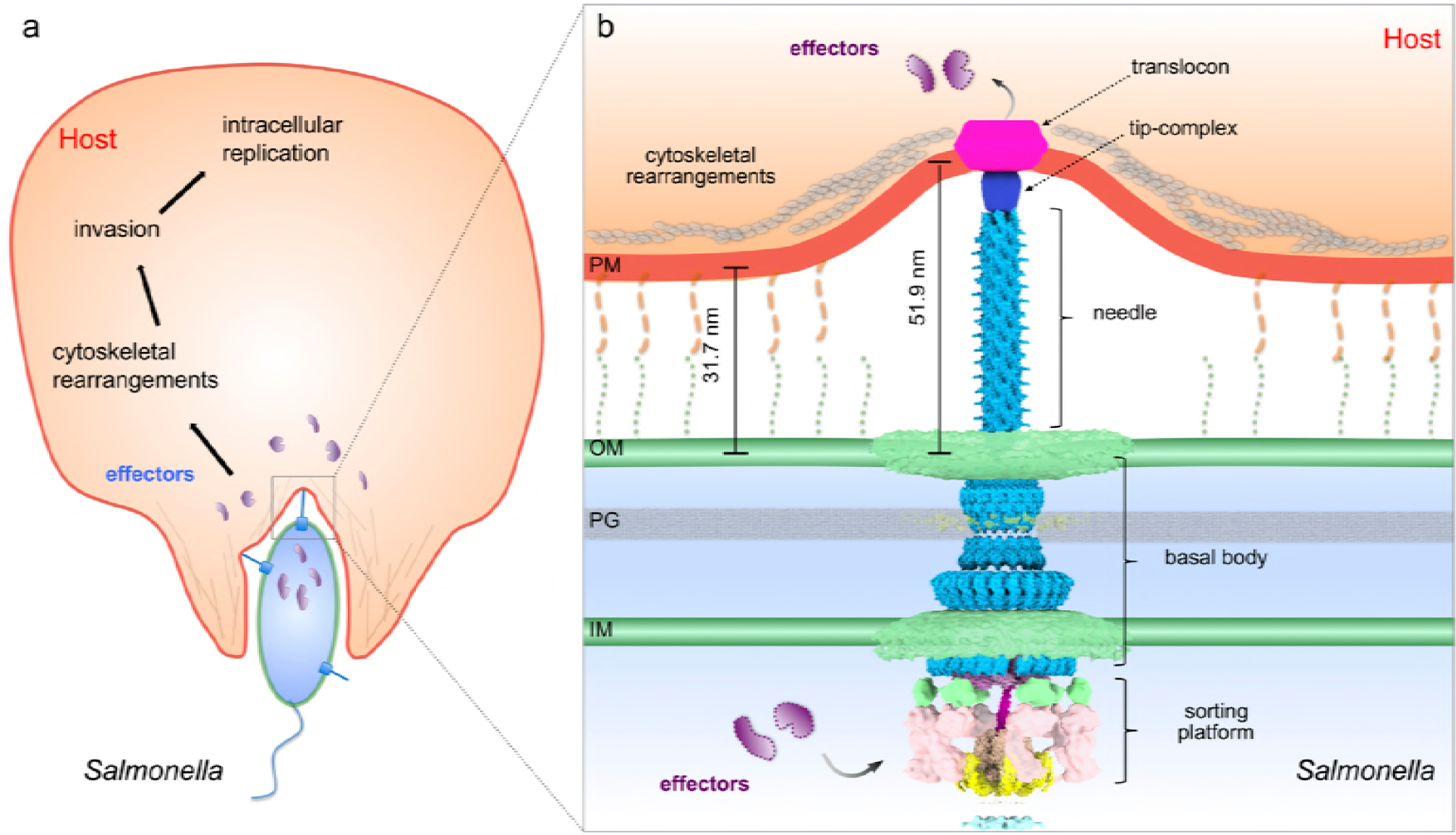
Model of the *S.* Typhimurium injectisome interacting with the host cell. (**a**) A schematic diagram of *S. Typhimurium* interacting with the host cell. (**b**) Molecular model of the T3SS injectisome at the *Salmonella*-host cell interface.

## DISCUSSION

We have presented here a high-resolution view of the interface between the *S*. Typhimurium T3SS injectisome and the target eukaryotic cell plasma membrane, which has provided details on the intimate attachment of this pathogen that precedes T3SS-mediated effector protein translocation. Notably, we observed a notable “bend” on the target cell PM in areas of the bacteria/PM interface surrounding the needle filament. These observations reflect the intimate attachment that is known to be required for optimal T3SS-mediated effector translocation that may result in the close apposition of the tip complex and the target cell PM. Importantly, we have been able to visualize a distinct density within the region of the target cell PM in close apposition to the needle tip of the T3SS injectisome. We present evidence that this density corresponds to the deployed T3SS translocon since this density was absent in the bacteria/PM interface of mutant bacteria that lack the translocon components. The dimensions of this structure (~13.5 nm in diameter, 8 nm in thickness) are much smaller than previous estimates (50-65 nm in diameter) obtained from the observation of EPEC translocons assembled from purified components on red blood cells (Ide et al, 2001). It is unlikely that these differences may reflect substantial differences between the dimensions of translocons from different T3SSs. It is possible that the observed differences may reflect differences in the experimental approaches used in the different studies. However, most likely these observations indicate fundamental differences in the translocon assembly mechanisms from purified components in comparison to translocon assembly during bacteria/target cell PM interactions. It is well established that the deployment of the translocon during bacterial infections is orchestrated by the needle filament tip complex of the T3SS injectisome. In the absence of the tip protein, the components of the translocon are very efficiently secreted but they are unable to form the translocon (Kaniga et al, 1995; Ménard et al, 1994). It is therefore possible that the insertion in the membrane of the purified translocon components in the absence of the tip protein may lead to a structure that is substantially different from the one that results from the interaction of bacteria with target cells.

Contrary to what has been previously proposed for the Chlamydia T3SS (Nans et al, 2015), we did not observe any obvious conformational changes in the injectisomes prior and post interaction with host cells. It is unlikely that these observations are an indication of fundamental differences between the T3SS injectisomes in different bacteria. Rather, the differences observed might reflect differences in the experimental approaches used in our studies, which resulted in a substantially higher resolution.

In summary, our studies have provided a close-up view of the interface between the T3SS injectisome and the target cell PM, which has resulted in the visualization of the deployed T3SS translocon complex. Importantly, given the highly conserved nature of the T3SSs among many Gram-negative bacteria, our studies have broad scientific implications and provide a paradigm for the study host-pathogen interactions in a greater detail.

## MATERIALS AND METHODS

***Bacterial strains.*** The minicell producing *S*. Typhimurium ∆*minD*, which is referred to in this study as wild-type, has been previously described (Carleton et al, 2013; Hu et al, 2017). Mutations in the genes encoding the translocases (*∆sipB, ∆sipC*) or tip complex *(∆sipD*) proteins where introduced in into the *∆minC S*. Typhimurium strain by allelic exchange as previously described (Lara-Tejero et al, 2011).

***Isolation of minicells.*** Minicell producing bacterial strains were grown overnight at 37 °C in LB containing 0.3M NaCl. Fresh cultures were prepared from a 1:100 dilution of the overnight culture and then grown at 37 °C to late log phase in the presence of ampicillin (200 μg/mL) and L-arabinose (0.1%) to induce the expression of regulatory protein HilA and thus increase the number of injectisomes partitioning to the minicells (Carleton et al, 2013). To enrich for minicells, the culture was centrifuged at 1,000 x g for 5 min to remove bacterial cells, and the supernatant fraction was further centrifuged at 20,000 x g for 20 min to collect the minicells. The minicell pellet was resuspended in Dulbecco’s Modified Eagles Medium (DMEM) prior to their application to cultured HeLa cells.

***HeLa cell culture on EM grid and infection.*** HeLa cells were cultured in DMEM supplemented with 10% fetal bovine serum and gentamicin (50 μg/ml). The day before plating, gold EM grids with 2/1 Quantifoil were placed in glass bottom MatTek dishes (facilitating fluorescence imaging and removal for cryo-preservation) and coated with 0.1 mg/ml poly-D-lysine overnight at 37°C. After rinsing the grids with sterile water, freshly trypsinized HeLa cells were plated on top of the pre-treated grids that were allowed to grow overnight at 37°C/5% CO2. To infect HeLa cells with *S*. Typhimurium minicells, grids with adherent HeLa cells were removed from the culture dish and minicells were directly applied to the grids.

***Vitrification and cryoEM sample preparation.*** At different time points after infection, the EM grids with HeLa cells and *S*. Typhimurium minicells were blotted with filter paper and vitrified in liquid ethane using a gravity-driven plunger apparatus as described (Hu et al, 2017; Hu et al, 2015).

***Cryo-ET data collection and reconstruction.*** The frozen-hydrated specimens were imaged with 300kV electron microscopes. 713 tomograms were acquired from single-axis tilt series at ~6 μm defocus with cumulative does of ∼80 e−/Å^2^ using Polara equipped with a field emission gun and a direct detection device (Gatan K2 Summit). 313 tomograms were acquired from single-axis tilt series at ~1 μm defocus with cumulative does of ∼50 e−/Å^2^ using Titan Krios equipped with a field emission gun, an energy filter, Volta phase plate, and a direct detection device (Gatan K2 Summit). The tomographic package SerialEM (Mastronarde, 2005) was utilized to collect 35 image stacks at a range of tilt angles between -51° and +51° for each data set. Each stack contained 10-15 images, which were first aligned using Motioncorr (Li et al, 2013) and were then assembled into the drift-corrected stacks by TOMOAUTO (Hu et al, 2015). The drift-corrected stacks were aligned and reconstructed by using marker-free alignment (Winkler & Taylor, 2006) or IMOD marker-dependent alignment (Kremer et al, 1996). In total, 1026 tomograms (3,600 ’ 3,600 ’ 400 pixels) were generated for detailed examination of the *Salmonella*-host interactions (Extended Data Table 3).

***Sub-tomogram analysis.*** Sub-tomogram analysis was accomplished as described previously (Hu et al, 2015) to analyze over 700 injectisomes extracted from 458 tomograms. Briefly, we first identified the injectisomes visually on each minicell. Two coordinates along the needle were used to estimate the initial orientation of each particle assembly. For initial analysis, 4 ’ 4 ’ 4 binned sub-tomograms (128 ’ 128 ’ 128 voxels) of the intact injectisome were used for alignment and averaging by using the tomographic package I3 (Winkler & Taylor, 2006; Winkler et al, 2009). Then multivariate statistical analysis and hierarchical ascendant classification were used to analyze the needle tip complex (Winkler et al, 2009).

***3-D visualization and molecular modeling.*** Outer membrane (OM) & inner membrane (IM) of *S*. Typhimurium, Plasma membrane (PM) of HeLa cells, actin filaments, and ribosomes were segmented using EMAN2 (Chen et al, 2017). UCSF Chimera (Pettersen et al, 2004) and UCSF ChimeraX (Goddard et al, 2018) were used to visualize the sub-tomogram average structures in 3-D and build atomic model of the T3SS injectisome. The atomic model was built as described briefly (Hu et al, 2017) except for the basal body, which we docked PDB-5TCR (Worrall et al, 2016) and PDB-3J1W (Bergeron et al, 2013). Video clips for the supplemental videos were generated using UCSF Chimera, UCSF Chimera X, and IMOD, and edited with iMovie.

***Distance measurement and statistical analysis.*** IMOD (3dmod Graph) was used to measure lengths (in pixels) of various features. Each measurement was recorded in MS Excel for statistical analysis: Mean, standard deviation, standard error of mean, and Welch’s t-test.

## ACKNOWLEDGEMENTS

This work was supported by Grants AI030492 (to J. E. G.) from the National Institute of Allergy and Infectious Diseases, and GM107629 from the National Institute of General Medicine (to J. L.).

